# Machine learning identifies SNPs predictive of advanced coronary artery calcium in ClinSeq® and Framingham Heart Study cohorts

**DOI:** 10.1101/102350

**Authors:** Cihan Oguz, Shurjo K Sen, Adam R Davis, Yi-Ping Fu, Christopher J O’Donnell, Gary H Gibbons

## Abstract

One goal of personalized medicine is leveraging the emerging tools of data science to guide medical decision-making. Achieving this using disparate data sources is most daunting for polygenic traits and requires systems level approaches. To this end, we employed random forests (RF) and neural networks (NN) for predictive modeling of coronary artery calcification (CAC), which is an intermediate end-phenotype of coronary artery disease (CAD). Model inputs were derived from advanced cases in the ClinSeq_®_ discovery cohort (n=16) and the FHS replication cohort (n=36) from 89^*th*^−99^*th*^ CAC score percentile range, and age-matching controls (ClinSeq® n=16, FHS n=36) with no detectable CAC (all subjects were Caucasian males). These inputs included clinical variables (CLIN), genotypes of 57 SNPs associated with CAC in past GWAS (SNP Set-1), and an alternative set of 56 SNPs (SNP Set-2) ranked highest in terms of their nominal correlation with advanced CAC state in the discovery cohort. Predictive performance was assessed by computing the areas under receiver operating characteristics curves (AUC). Within the discovery cohort, RF models generated AUC values of 0.69 with CLIN, 0.72 with SNP Set-1, and 0.77 with their combination. In the replication cohort, SNP Set-1 was again more predictive (AUC=0.78) than CLIN (AUC=0.61), but also more predictive than the combination (AUC=0.75). In contrast, in both cohorts, SNP Set-2 generated enhanced predictive performance with or without CLIN (AUC> 0.8). Using the 21 SNPs of SNP Set-2 that produced optimal predictive performance in both cohorts, we developed NN models trained with ClinSeq® data and tested with FHS data and replicated the high predictive accuracy (AUC>0.8) with several topologies, thereby identifying several potential susceptibility loci for advanced CAD. Several CAD-related biological processes were found to be enriched in the network of genes constructed from these loci. In both cohorts, SNP Set-1 derived from past CAC GWAS yielded lower performance than SNP Set-2 derived from “extreme” CAC cases within the discovery cohort. Machine learning tools hold promise for surpassing the capacity of conventional GWAS-based approaches for creating predictive models utilizing the complex interactions between disease predictors intrinsic to the pathogenesis of polygenic disorders.

## BACKGROUND

Informed medical decision making through the effective use of clinical and genomic data is one of the promising elements of personalized precision medicine (Ginsburg and Willard, 2009) in which predictive models enable the systematic assessment of alternative treatment approaches taking into account the genomic variability among different patients (Völzke et al., 2013). Predictive models not only play a pivotal role in utilizing the genomic data for generating predictions regarding the disease risk and state (Cui and Lincoln, 2015; Jiang et al., 2012; Khorana et al., 2008; Hood et al., 2004; Bellazzi and Zupan, 2008; Nevins et al., 2003; West et al., 2006), but they may also generate biological insights into the mechanisms behind complex diseases (Lee et al., 2013), such as coronary artery disease (CAD) that claims the lives of millions of people globally as the leading cause of death (Santulli, 2013). In CAD, the arteries of the heart, which supply oxygen rich blood to the cardiac muscle, lose their ability to function properly due to atherosclerosis. CAD is a multifactorial disease (Poulter, 1999; Schwartz et al., 2012) that has been associated with a large number of clinical and demographic variables, and major risk factors such as high blood pressure, high levels of blood lipids, smoking and diabetes. Our main focus in this study, namely coronary artery calcification (CAC), is an intermediate end-phenotype of CAD (McClelland et al., 2014) and a strong predictor of cardiac events including myocardial infarction (MI) (Forster and Isserow, 2005; Williams et al., 2014; Liu et al., 2013; Wayhs et al., 2002; Budoff et al., 2009, 2013). This predictive feature of CAC has been a major driving force behind research on its statistical characterization as an intermediate phenotype for CAD in recent years (Sun et al., 2008; McGeachie et al., 2009; Natarajan et al., 2012).

The key mechanism behind coronary artery calcification is the phenotypic modulation of vascular cells into a mineralized extracellular matrix (ECM) (Johnson et al., 2006). This modulation is triggered by stimuli including oxidative stress, increased rate of cell death (Proudfoot et al., 2000; Kim, 1994), and high levels of inflammatory markers (Rutsch et al., 2011; Johnson et al., 2006). The genetics behind coronary calcium deposition is fairly complex, which is not surprising given that it is a commonly observed phenomenon (common disease phenotypes are typically multigenic (Swan, 2010)). Several important genes involved in vascular calcification have been previously identified through mouse model studies (Nitschke and Rutsch, 2014; Rutsch et al., 2011), studies on rare human diseases that lead to excessive calcification (Rutsch et al., 2011), as well as through elucidation of the links between bone mineralization and CAC (Marulanda et al., 2014). Several genome-wide association studies (GWAS) have also previously focused on CAC (Ferguson et al., 2013; Wojczynski et al., 2013; van Setten et al., 2013; O’Donnell et al., 2007, 2011; Polfus et al., 2013). Some of the human genomic loci associated with CAC through GWAS are *9p21*, *PHACTR*, and *PCSK9*, all of which have been also linked to CAD and MI (van Setten et al., 2013; Kathiresan et al., 2009; Dubuc et al., 2010). Several past studies have combined clinical variables and genotype data in order to improve predictions for CAD. Some examples include implementation of Cox regression models (Morrison et al., 2007; Brautbar et al., 2012; Kathiresan et al., 2008) and the use of allele counting, logistic regression, and support vector machines in (Davies et al., 2010). Even though multiple studies showed statistically significant improvements in predicting CAD by combining traditional risk factors with SNPs linked to CAD in past GWAS, the reported improvements have been at best incremental (Ioannidis, 2009). Similar results have been compiled in a recent review (Liao and Tsai, 2013) for type 2 diabetes (a strong risk factor for CAD) where marginal improvements were observed in some studies.

Recently, there has been increasing interest in the application of machine learning methods for predicting disease phenotypes by utilizing genomic features (Goldstein et al., 2016). These methods provide increased ability for integrating disparate sources of data while utilizing interactions (both linear and nonlinear) between genomic features (e.g., gene-gene interactions) unlike conventional regression approaches (Chen and Ishwaran, 2012). Machine learning methods also eliminate a major limitation of GWAS, which is the need for multiple testing correction required in statistical association tests that treat each predictor separately, while also avoiding biases that could originate from model misspecification since machine learning typically aims at identifying model structures that are optimal for the training data (Li et al., 2015).

In this study, we utilized machine learning tools for predictive modeling of advanced coronary calcification among Caucasian males by integrating clinical variables and genotype data. Our study focused on Caucasian males due to higher coronary calcium scores observed among men compared to women (Raggi et al., 2008; Maas and Appelman, 2010), as well as higher prevalence of coronary calciumamong white Americans compared to black Americans (Lee et al., 2003). Using random forest modeling, which is a decision tree based machine learning method (Breiman, 2001) established as an effective tool for addressing the complexity of modelling with genomic data (Sun, 2009; Yang et al., 2010b; Dietterich, 2000), we first tested the collective ability of a set of SNPs derived from previous GWAS on CAC (SNP Set-1) in predicting advanced CAC with data from the ClinSeq® study (Biesecker et al., 2009) previously published in (Sen et al., 2014b,a). Upon deriving an alternative SNP set (SNP Set-2) and comparing its predictive ability to SNP Set-1 within the ClinSeq® discovery cohort with and without clinical data, we used data from the Framingham Heart Study (FHS) to test whether we could replicate the observed predictive patterns. Then, in order to identify a set of potential susceptibility loci for advanced CAD pathogenesis, we derived the subset of SNPs in SNP Set-2 that led to optimal predictive performance in both cohorts. Using this subset of SNPs, we developed neural network models trained with data from the ClinSeq® discovery cohort and tested with data from the FHS replication cohort under a wide range of network topologies, assessed the predictive performances of these models, and identified the biological processes enriched in the network of genes constructed from the predictive loci.

## METHODS

### Overview of the computational analysis

As illustrated in Figure 1, the overall strategy of our analysis was to initially use only clinical data for predicting advanced CAC in a discovery cohort, then to combine clinical data with a GWAS-based set of SNPs to test for improved predictive performance in a discovery cohort. We also aimed to derive an alternative set of SNPs that are collectively more predictive in this discovery cohort and to test if the observed predictive patterns were replicable with or without clinical data in an independent replication cohort. In order to achieve these objectives, we took the following steps as shown in Figure 2. First, we developed random forest models that predict advanced CAC within the ClinSeq® cohort that served as our “discovery cohort” using traditional risk factors (or clinical variables) and a set of GWAS-identified SNPs (or “SNP Set-1”) previously associated with coronary calcium. We assessed the predictive performance by using only clinical data (to establish a baseline performance) or genotype data, as well as their combination. We then derived a second set of SNPs (or “SNP Set-2”) as an alternative to SNP Set-1 using data from the discovery cohort utilizing a selection criterion based on the nominal correlations between SNP genotypes and the advanced CAC state.

**Figure 1.**
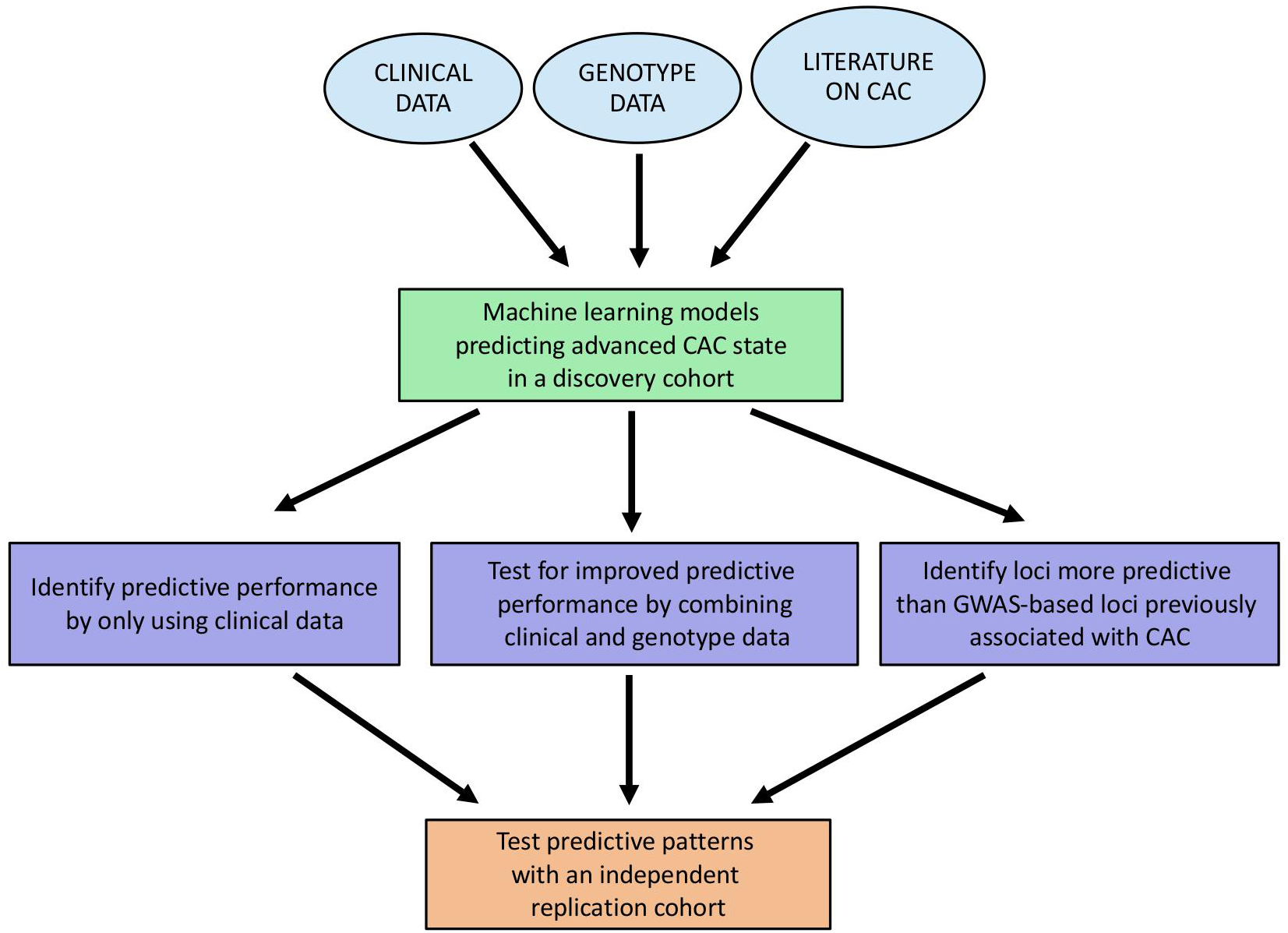
Overall strategy of the analysis.

**Figure 2.**
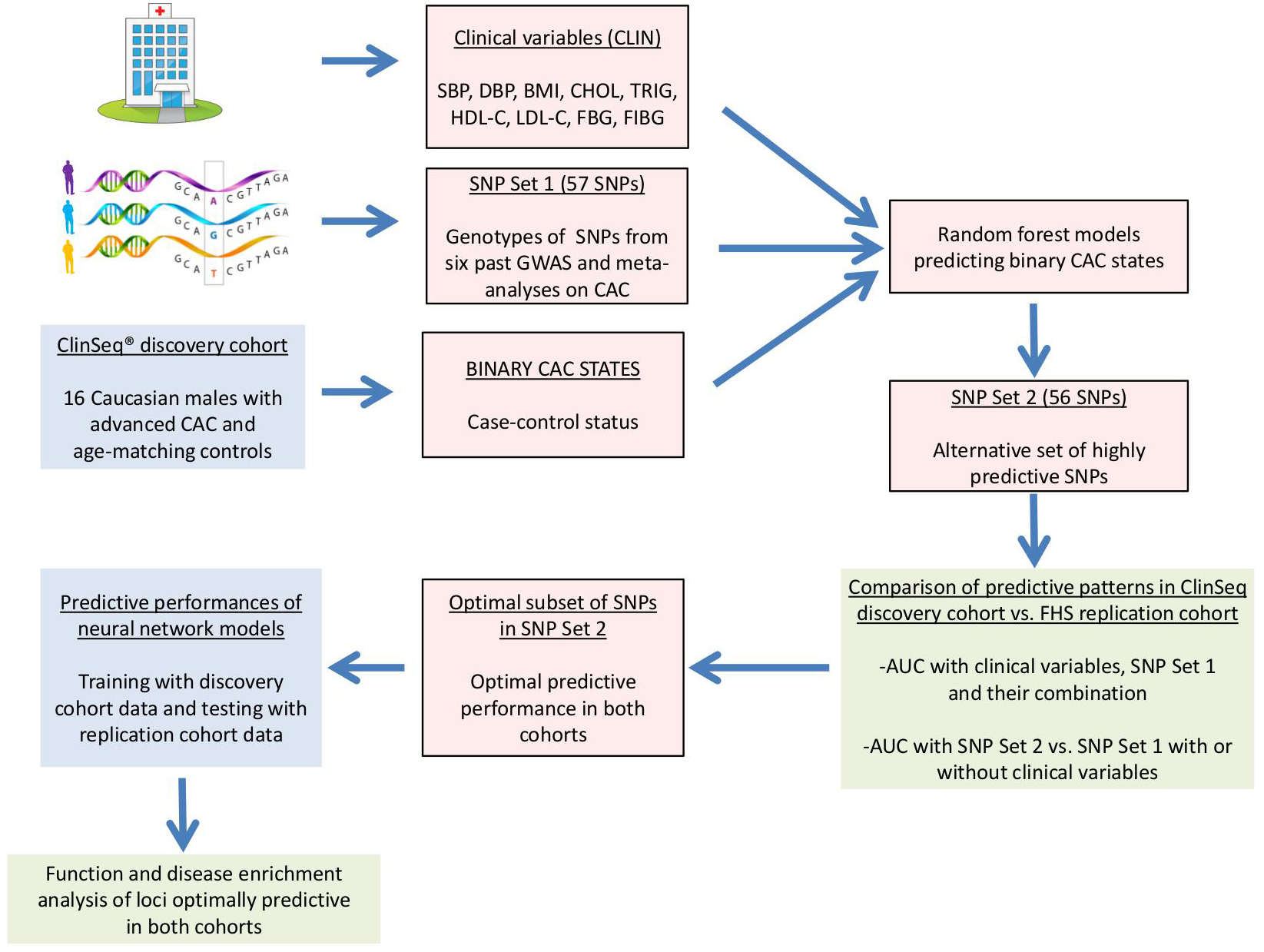
Schematic of the modeling approach.

Upon comparing the random forest based predictive patterns generated by the clinical variables, SNP Set-1, and SNP Set-2 in the ClinSeq® discovery cohort and the FHS replication cohort, we identified the subset of SNPs in the more predictive set that generated optimal performance in random forest models of both cohorts. We trained neural network models with the genotypes of these SNPs among all ClinSeq® subjects and tested with the genotypes of the same SNPs among all FHS subjects with the aim of obtaining high predictive accuracy values under a wide range of neural network topologies. We then utilized GeneMANIA (Warde-Farley et al., 2010; Zuberi et al., 2013; Montojo et al., 2014) to create a functional interaction network composed of genes on which this subset of SNPs was located, as well as additional genes known to be most closely related to these genes. GeneMANIA uses linear regression to maximize the connectivity between the genes within the network while minimizing the interactions with the genes that are excluded. Two types of links between gene pairs were found to be present in this network: co-expression (correlated expression levels) and genetic interactions (effects of a gene perturbation can be changed by a second perturbed gene). Gene Expression Omnibus (GEO) and BioGRID are the main sources of co-expression and genetic interaction datasets, respectively in the GeneMANIA database. Finally, using the list of genes within this network derived by GeneMANIA, we performed function and disease enrichment analysis to demonstrate the relevance of these advanced coronary calcium susceptibility loci to cardiovascular disease based on the existing knowledge in the literature.

### Coronary calcification scores and binary CAC states

The models we developed in this study aimed at predicting the binary case-control statuses of Caucasian male patients. Hence, we first transformed the CAC scores (measured by Agatston method (Agatston et al., 1990)) of the 32 Caucasian male subjects from the ClinSeq® study that formed our discovery cohort (data previously published in (Sen et al., 2014a, b)) into binary CAC states. 16 control subjects in this cohort had zero CAC scores corresponding to state “0”, whereas the 16 age-matching cases had high CAC scores (ranging between 500 and 4400) corresponding to state “1”. These binary case-control states served as the true class labels and were later used for training and testing of the developed classification models. Based on the Multi-Ethnic Study of Atherosclerosis (MESA) cohort standards (McClelland et al., 2006; Dat, 2015), a percentile value for each case was computed using the online MESA calculator (Dat, 2015) that takes age, gender, race and CAC score as its inputs. The case subjects in the ClinSeq® discovery cohort, two of which were diabetic, fell within the 89^*th*^-99^*th*^ CAC score percentile range.

Figure 1. Overall strategy of the analysis.

The replication cohort from FHS comprised of 36 controls and 36 age-matching Caucasian male case subjects (including three diabetic cases) also within the 89^*th*^-99^*th*^ CAC score percentile range. Additional 122 cases from FHS within 29^*th*^-88^*th*^ CAC score range were split into two distinct sets of 61 cases within 29^*th*^-68^*th*^ and 69^*th*^-88^*th*^ percentile ranges and were age-matched with two sets of 61 controls. These two equal-sized subcohorts were then used to test whether the predictive patterns generated by the discovery (ClinSeq®) and replication (FHS) cohorts were specific to the 89^*th*^-99^*th*^ percentile CAC score range and not replicable with lower levels of coronary calcium. Two classes of model variables were used in this study as predictors of coronary calcium, namely clinical variables and genotypic variables, as described below.

### Clinical variables

Nine clinical variables available from all subjects in both cohorts were utilized as predictors of coronary calcium. These variables included body mass index (BMI), cholesterol levels (LDL, HDL, and total cholesterol), triglycerides, blood pressure (systolic and diastolic), fasting blood glucose level, and fibrinogen. All subjects were non-smoker Caucasian males in both ClinSeq® and FHS cohorts. The detailed description of each clinical variable is given in Table S1, whereas the mean and standard deviation values among cases vs. controls, along with their p-values are listed in Tables S2 and S3 for ClinSeq® and FHS cohorts, respectively.

### Genotypic variables

For the ClinSeq® cohort, SNP genotyping was performed using the HumanOmni2.5 Illumina BeadChip arrays. Genotyping was carried out with in accordance with the Illumina Infinium assay protocol. In brief, this involved amplification of DNA by WGA, hybridization of the WGA product to the BeadArray, an array-based enzymatic reaction that extends the captured SNP targets by incorporating biotin-labeled dNTP nucleotides into the appropriate allele specific probe, and, finally, detection and signal amplification to read the incorporated labels. The BeadChips were scanned using the Illumina iScan system and processed with the GenomeStudio v2011.1 Genotyping module. The BeadChips consist of specific 50-mer oligonucleotide probe arrays at an average of 30-fold redundancy. The design of the HumanOmni2.5 BeadChips incorporates around 2.5 million markers. GenomeStudio output files were processed using a custom Perl script to derive the nucleotides at each SNP position for each subject.

For the FHS cohort, genotyping data was compiled from three resources. More than 276,000 variants from the Illumina Infinium Human Exome Array v1.0 was genotyped and jointly called as part of the Cohorts for Heart and Aging Research in Genomic Epidemiology (CHARGE) Consortium (PMID 23874508). The Framingham SNP Health Association Resource (SHARe) project (Dat, 2016a) used the Affymetrix 500K mapping array and the Affymetrix 50K supplemental gene focused array resulted in 503,551 SNPs with successful call rate >95% and Hardy-Weinberg equilibrium (HWE) P>1.0E-6. Additional genotype imputation was conducted based on this SHARe data using Minimac with reference panel from the 1000 Genomes Project (Version Phase 1 integrated release v3, April 2012, all population). Best-guessed genotypes with imputation quality >0.3 were used for markers that were not available from the first two actual genotyping platforms.

We compiled a set of 57 SNPs (listed in Table S4) that were associated with coronary calcium in previous GWAS (Ferguson et al., 2013; Wojczynski et al., 2013; O’Donnell et al., 2011, 2007; Polfus et al., 2013; van Setten et al., 2013) and named this set “SNP Set-1”. From the the ClinSeq® genotype data, we also generated a second set of SNPs (“SNP Set-2”), approximately the same size (56) as the “SNP Set-1”, by using a genotype-phenotype correlation criterion as listed in Table S5. Genotypes of the 113 biallelic SNPs in both SNP sets were coded as 0 or 2 (homozygous for either allele) or 1 (heterozygous) using the same reference alleles in both ClinSeq® and FHS cohorts.

### Predictive modeling using random forests and neural networks

We implemented the random forest classification method using the Statistics and Machine Learning Toolbox^™^ of Matlab® (MATLAB, 2013) for predicting the binary CAC state. Predictive accuracy is computed by generating ROC curves (true positive rate vs. the false positive rate obtained using several classifier output thresholds) and quantifying the areas under these curves (AUC). Due to the randomized nature of the classification method, we performed 100 runs (per set of features or model inputs) and reported the mean AUC (AUC distributions were normal based on the Anderson-Darling tests (Stephens, 1974)) and its p-value that is derived empirically (Ojala and Garriga, 2010; Sun et al., 2008) by performing 1000 runs with randomly permuted case-control statuses and computing the fraction of AUC values below the mean AUC value generated when the case-control statuses are not permuted (i.e., the actual data), an approach commonly used for computing the statistical significance of AUC in ROC-based predictive modeling studies. Per decision tree, approximately two-thirds of the data (this ratio varied up to 15% among different runs) is retained to be used for model training, whereas the remaining data is used for model testing. These test samples are referred to as “out-of-bag” (OOB) samples, whereas the training samples are expanded by bootstrapping (Efron, 1979) (or sampling with replacement) up to the sample size of the original data (Dasgupta et al., 2011) prior to model training. Classification of the test samples are based on the complete ensemble of trees (a total of 100 trees) with a voting scheme. For example, a test sample is predicted to be “CAC positive” if the number of trees that predict “State 1” is higher than the ones that predict “State 0”. Predictive importance is computed for each input variable by permuting its values corresponding to the test subjects and finding the change in the prediction error (or the fraction of incorrectly classified subjects). One error value is computed for each tree and the ratio of the average value of this change is divided by the standard deviation. Features are ranked with respect to this ratio (i.e., features that are stronger predictors have higher values of this ratio compared to the weaker predictors). After ranking all features in each distinct feature set (e.g., all clinical variables), we decreased the number of features gradually by leaving out weaker predictors to identify the optimal predictive performance and the corresponding optimal set of features. We repeated this procedure to compare the predictive performances of models trained and tested by combining clinical and genotype data, as well as using each layer data in isolation. By identifying the predictive SNPs in GWAS-based SNP Set-1 and the alternative SNP Set-2, we were also able to compare the cumulative predictive importance scores from SNP-risk factor and SNP-CAD related phenotype associations (tied to each SNP set) based on past studies. The predictive patterns generated by data from the ClinSeq® discovery cohort were also compared with the patterns generated by the independent FHS replication cohort. Finally, random forest models were also used to identify a subset of SNPs in SNP Set-2 that generated the optimal predictive performance in both ClinSeq® and FHS cohorts.

Upon identifying the subset of SNPs in SNP Set-2 that generate random forest models with optimal performance in both cohorts, we implemented a neural-network-based classification approach using the Neural Network Toolbox^™^ of Matlab® (MATLAB, 2013). We trained three-layer feedforward networks using backpropagation (Fausett, 1994) with sigmoid transfer functions in two hidden layers and a linear transfer function in the output layer. In both hidden layers, the number of nodes was varied from one to 20 with increments of one, thereby leading to a total of 400 network configurations individually used for training and testing. In short, the inputs into each network layer (initial input is the genotype data) are weighted and the sum of the weighted inputs transformed by the transfer functions of the hidden layers are used to generate model outputs (or the case/control status) (Mehrotra et al., 1997). We trained all network configurations with the genotypes of the optimal subset of SNPs within SNP Set-2 from all subjects in the ClinSeq® discovery cohort (approximately 20% of these samples include the “validation” samples used for minimizing overfitting during training with the remaining 80% of the samples) and subsequently performed model testing with the genotype data from all subjects in the FHS replication cohort. Predictive accuracy was once again assessed with ROC curves. For each neural network configuration, we computed the median AUC value (AUC distributions were non-normal based on the Anderson-Darling tests (Stephens, 1974)) among 100 independent runs and empirically derived the p-value as the fraction of AUC values from 1000 runs with randomly permuted case-control statuses below the median AUC value obtained when the case-control statuses are not permuted (i.e., actual data).

## RESULTS

### Models built with clinical variables and GWAS-based SNP Set-1

We first built random forest models using all of the nine clinical variables from the ClinSeq® discovery cohort and identified that three of them had positive predictive importance values as listed in Table These predictors included HDL Cholesterol (a major risk factor for coronary calcium (Allison and Wright, 2004; Parhami et al., 2002)), systolic blood pressure, and fibrinogen that has been previously associated with vascular calcification (Bielak et al., 2000; Rodrigues et al., 2010) as a critical parameter for inflammation (Davalos and Akassoglou, 2012) and atherosclerosis (Smith, 1986). Within the FHS replication cohort, five clinical variables including total cholesterol, systolic and diastolic blood pressure, fibrinogen and fasting blood glucose (a glycemic trait previously associated with high coronary calcium levels (Schurgin et al., 2001)) had positive predictive importance values. The aggregate of the clinical variables with predictive power in the discovery and replication cohorts formed a combination of lipid and glycemic traits with a blood coagulation trait reflecting a “metabolic syndrome” picture (Eckel et al., 2005; Nieuwdorp et al., 2005). As we varied the number of predictors between one to nine, the optimal AUC values were 0.69 (p-value=0.015) and 0.61 (p-value=0.080) for ClinSeq® and FHS cohorts, respectively (Figure 3A). These AUC values were within the range of 0.60-0.85, which is the previously reported AUC range compiled from 79 studies predicting CAD or cardiac events based on the Framingham risk score (FRS) (Tzoulaki et al., 2009), despite our inability to use age and gender in predicting advanced CAC due to the design of our study.

**Table 1.**
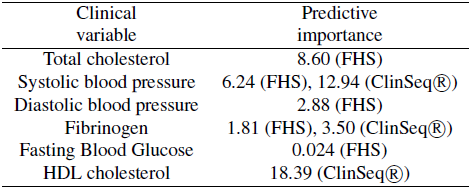
Predictive importance values of clinical variables in ClinSeq® and FHS cohorts. Only the instances with positive predictive importance are reported.

| Clinical variable | Predictive importance |
| --- | --- |
| Total cholesterol | 8.60 (FHS) |
| Systolic blood pressure | 6.24 (FHS), 12.94 (ClinSeq®) |
| Diastolic blood pressure | 2.88 (FHS) |
| Fibrinogen | 1.81 (FHS), 3.50 (ClinSeq®) |
| Fasting Blood Glucose | 0.024 (FHS) |
| HDL cholesterol | 18.39 (ClinSeq®) |

**Figure 3.**
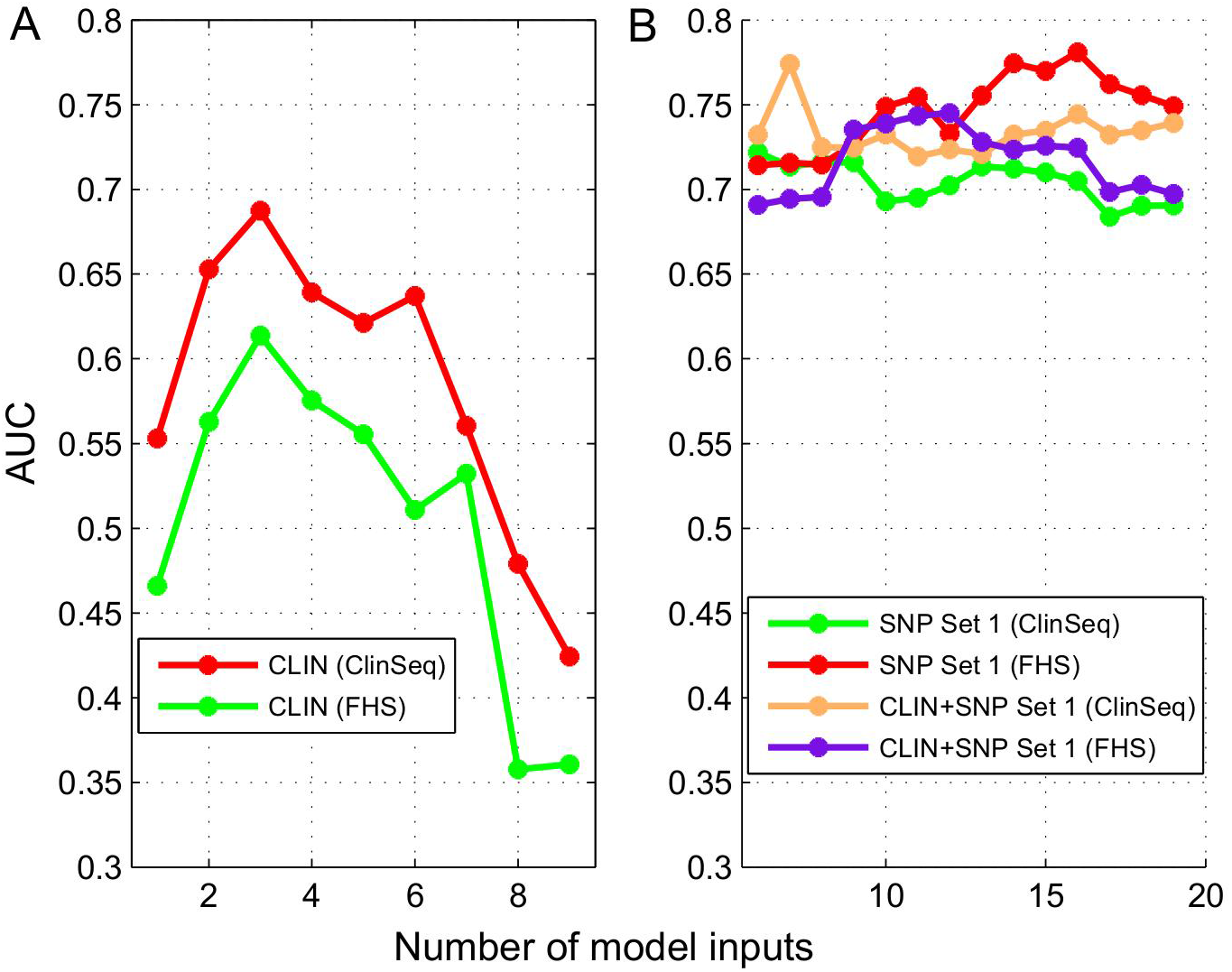
Predictive performance vs. number of predictors in ClinSeq®and FHS cohorts with onlyclinical variables in (A) and the combination of clinical variables and SNPs from SNP Set-1 in (B).

We next built random forest models for the ClinSeq® discovery cohort using the genotypes of the 57 SNPs in “SNP Set-1” as model inputs and identified 17 SNPs with positive predictive importance. In past GWAS, 11 of the 17 predictive SNPs have previously been associated with 18 CAD risk factors forming 28 SNP-risk factor pairs (Table S6), whereas six of them have been linked to CAD, MI, stroke, and aortic valve calcium (Table S7). For a detailed discussion of these associations and loci (including PCSK9 and 9p21), we refer the reader to Supplementary Text (Section 1).

To compare the predictive patterns generated by the discovery and replication cohorts based on the SNP Set-1 genotype data, we next developed random forest models for the FHS replication cohort and identified 19 SNPs among SNP Set-1 with positive predictive importance in this cohort. Figure 3B shows the AUC ranges as 0.68-0.72 and 0.71-0.78 for the ClinSeq® and FHS cohorts with the top 6-19 predictors (without clinical variables), respectively. Despite a small degree of overlap between these two ranges, only five of the 17 predictive SNPs (29%) from the ClinSeq® discovery data were replicated with the FHS data and only one of these five SNPs had predictive importance values in FHS and ClinSeq® data sets with magnitudes within 10% of each other (difference divided by the maximum value) pointing to a fairly low degree of replication between the two cohorts when only the GWAS-based SNP Set-1 is used for predicting advanced CAC. In order the test whether the combination of the two groups of predictors (nine clinical variables and SNP Set-1) resulted in improved predictive performance, we merged the two groups of model inputs with the ClinSeq® discovery data set. We observed a significant improvement in the AUC range from 0.68-0.72 (only SNP Set-1) to 0.72-0.77 (combined set of inputs) with the top 6-19 predictors as shown in Figure 3B. In contrast, when we used the FHS replication data set in the same way, AUC range declined from 0.71-0.78 to 0.69-0.75. Hence, the improvement of predictive accuracy we observed within the ClinSeq® discovery cohort, by adding clinical variables to SNP Set-1, was not observed with the FHS replication cohort (Table 2). This outcome pointed out another limitation of the past GWAS-based SNP Set-1 since the improvement of accuracy observed in the discovery cohort by combining clinical variables and SNP Set-1 as model inputs was not replicated in the FHS cohort.

**Table 2.** Predictive performances of RF models (quantified by the mean ± standard deviation values ofAUC) trained and tested with different predictor sets in the ClinSeq® and FHS cohort data. AUC distributions were normal based on the Anderson-Darling tests (Stephens, 1974). “CLIN” corresponds to the nine clinical variables listed in Table S1 (all variables except age and gender).

| Predictors | Optimal # markers<br>(ClinSeq®, FHS) | Optimal AUC<br>(ClinSeq®, FHS) | p-value<br>(ClinSeq®, FHS) |
| --- | --- | --- | --- |
| CLIN | 3, 3 | 0.69±0.02, 0.61±0.02 | 0.015, 0.080 |
| SNP Set-1 | 6, 16 | 0.72±0.02, 0.78±0.02 | 0.021, <0.001 |
| CLIN+SNP Set-1 | 7, 12 | 0.77±0.03, 0.75±0.02 | 0.013, <0.001 |
| SNP Set-2 | 21, 21 | 0.99±0.01, 0.85±0.02 | <0.001, <0.001 |
| CLIN+SNP Set-2 | 21, 18 | 0.99±0.01, 0.83±0.01 | <0.001, <0.001 |

### Selection of SNP Set-2 based on genotype-phenotype correlation within the ClinSeq® discovery cohort

Previous GWAS and meta-analyses studies on CAC focused on the presence of coronary calcium (including low levels of CAC), rather than its extreme levels. Since our discovery and replication cohorts both included cases with CAC scores within 89^*th*^-99^*th*^ percentile range, we next targeted the ClinSeq® discovery cohort genotype data to identify SNPs highly predictive of advanced CAC in order to utilize the advantages provided by random forest models (ability to identify optimal model structure for training data while utilizing interactions between SNPs without multiple testing penalty) over GWAS and conventional regression approaches. Expecting a correlation between the SNP genotype and the binary advanced CAC state (healthy control vs. 89^*th*^-99^*th*^ percentile CAC score range) for such predictive SNPs, we used a genotype-phenotype correlation criterion to identify an additional SNP set with approximately the same size as the SNP Set-1 from the ClinSeq® discovery cohort data. First, we verified the rationale behind the implemented genotype-phenotype correlation criterion. As shown in Figure 4, the predictive importance values of the SNPs in SNP Set-1 and the correlation between each SNP’s genotype and the case-control statuses of our subjects were highly correlated with each other. Here, the Pearson-based correlation coefficient was computed as 0.73 with a p-value of 2.61E-10 estimated by a two-tailed t-test. In simple terms, for any SNP within SNP Set-1, a higher correlation between the genotype and the case-control statuses of the 32 subjects led to a higher predictive importance value. Using this rationale, we identified an alternative “SNP Set-2” (56 SNPs not associated with CAC in past studies) whose genotypes had the highest correlation values with the case-control status. Within the ClinSeq® discovery cohort, the range of genotype-phenotype correlation among the SNPs in SNP Set-2 was 0.63-0.73, whereas the same range was 0-0.51 among the SNPs in SNP Set-1. Hence, there was no overlap between the two sets of SNPs.

**Figure 4.**
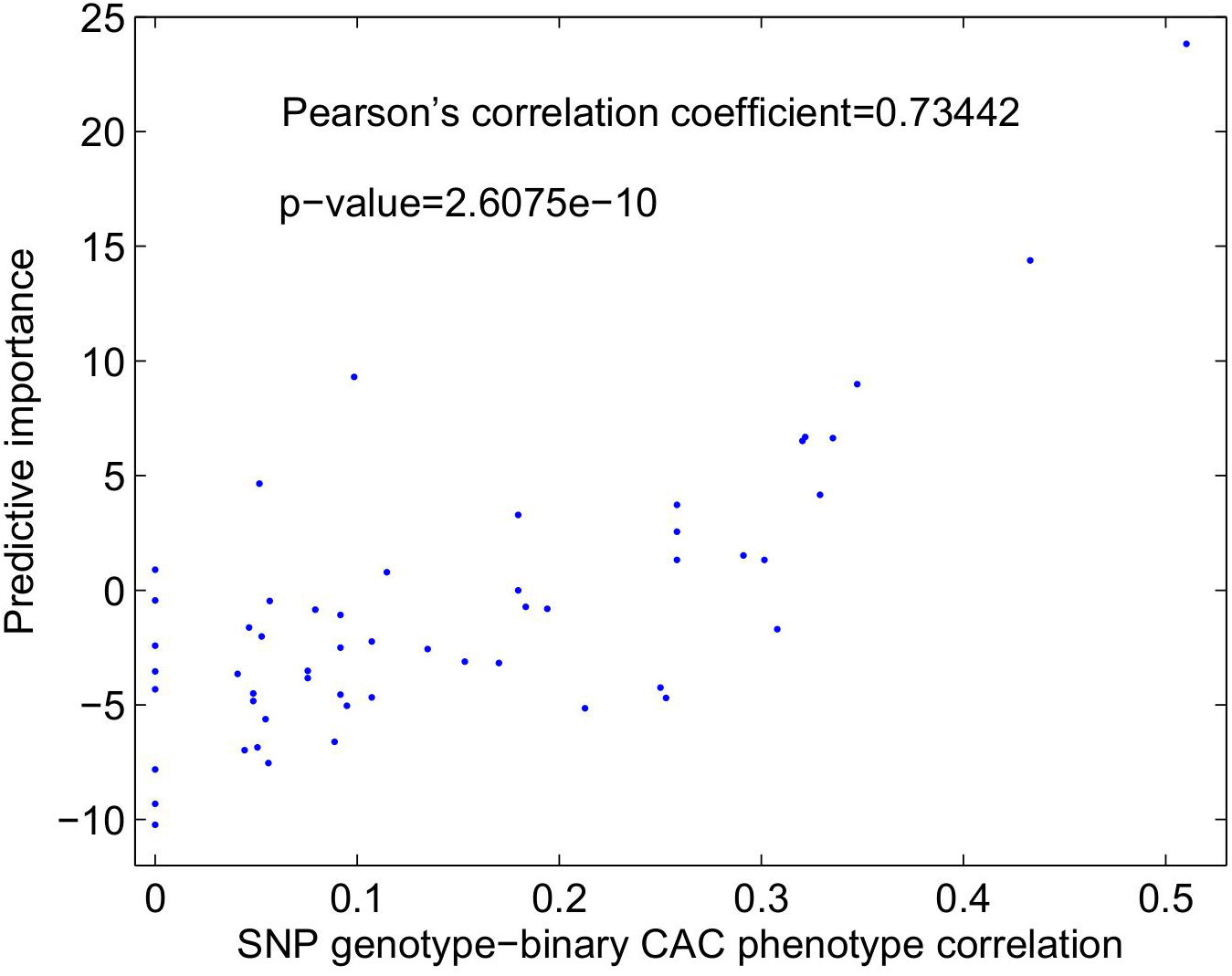
Predictive importance of the SNPs in SNP Set-1 vs. SNP genotype-binary CAC phenotype within the ClinSeq® discovery cohort. The strong correlation is indicated by the high Pearson’s correlation coefficient value and its corresponding p-value.

Upon incorporating the genotypes of SNP Set-2 within the ClinSeq® discovery cohort into random forest models, the AUC value turned out to be 0.9975, thereby verifying the superb ability of this set of markers. As shown in Table S8, 42 of these 56 predictive SNPs have been previously associated with a total of 18 risk factors, whereas the total number of SNP-risk factor pairs was 86 with many individual SNPs being associated with multiple risk factors. This was in contrast to only 11 of the 17 predictive SNPs in SNP Set-1 that were associated with a total of 18 risk factors forming 28 SNP-risk factor pairs. In addition, the susceptibility score, which is computed as the cumulative predictive importance values of SNPs tied to CAD risk factors in previous GWAS, increased from 446 to 1229 aligning with the improved predictive accuracy from 0.72 (maximum AUC in Figure 3B for SNP Set-1 in the ClinSeq® discovery cohort) to ≈1.00 as we moved between the two sets of predictors, namely SNP Set-1 and SNP Set-2. Table S9 shows that ten of the predictive SNPs in Set-2 that have been associated with stroke and aortic valve calcium in past GWAS, a trend we also observe with three SNPs in SNP Set-1 (Table S7). Two of the predictive SNPs in Table S9 have also been linked to mitral annular calcium, another disease phenotype related to coronary artery calcification along with aortic valve calcification, all of which are considered as common elements of atherosclerosis (Atar et al., 2003). The aggregate of the associations listed in Tables S8 and S9 suggests that the highly predictive SNPs identified from the ClinSeq® discovery cohort data (or SNP Set-2) could be potential susceptibility loci for advanced CAC.

### Comparing predictive performance of SNP Set-2 using FHS and ClinSeq® data sets

In order to test whether the higher predictive performance of SNP Set-2 over the past GWAS-based SNP Set-1 was replicated in the FHS cohort, we trained and tested random forest models using the genotypes of SNP Set-2 from the replication cohort. We identified that the positive predictive importance values of 30 of the 56 predictive SNPs (54%) were replicated. The predictive importance values of five SNPs in the two data sets were within 10% of each other, whereas nine SNPs had values within 20% of each other. We also observed common patterns between the discovery and replication cohorts in terms of the predictive importance based rankings of the 30 SNPs with positive predictive importance in both cohorts. Nine of the top 18 SNPs overlapped between the two cohorts, whereas the top two SNPs (rs243170 and rs243172, both on *FOXN3*) were the same in both cohorts. *FOXN3* is involved in transcription regulation at the cellular level and the G2/M phase of the cell cycle as a checkpoint suppressor. *FOXN3* has also been linked to fasting blood glucose in past GWAS (Manning et al., 2012) and in a recent study through its overexpression in human liver cells and zebrafish (Karanth et al., 2016).

Top 9-28 of the 30 SNPs with positive predictive importance generated AUC ranges of 0.80-0.85 and 0.96-0.99 in the replication and discovery cohorts, respectively. Based on these results, and given that the SNP Set-1 failed to reach an AUC value of 0.8 in both cohorts even with the optimal number of SNPs, we concluded that the higher predictive accuracy of SNP Set-2 over SNP Set-1 in the ClinSeq® discovery cohort was replicated in the FHS replication cohort. Combining the clinical variables and SNP Set-2 did not improve the predictive performance, consistently in both cohorts. In fact, there was a slight decline in the optimal AUC from 0.85 to 0.83 with the top 12-22 predictors in the FHS cohort, whereas no change in the optimal AUC was observed in the ClinSeq® cohort with the combination of clinical variables and SNP Set-2.

One potential explanation of the higher predictive performance of SNP Set-2 over SNP Set-1 in both cohorts is the broad CAC levels that were focused on past GWAS and meta-analyses (instead of highly advanced CAC) in order to reach adequate statistical power. Given that SNP Set-2 was derived from cases with extreme levels of CAC, it remained to be determined whether the predictive power of SNP Set-2 was specific to this extreme phenotype or whether it could be generalized to a broader range of CAC levels. Hence, we tested the collective predictive performance of the 30 SNPs in SNP Set-2 that had positive predictive power in both cohorts with genotype data from cases with lower levels of CAC. To achieve this, we used the genotype data of 122 cases from FHS within 29^*th*^-88^*th*^ percentile CAC score range. Among the 61 cases within the 29^*th*^-68^*th*^ percentile range and the 61 age-matching controls, top 9-28 markers generated an AUC range of 0.62-0.66, whereas only 20 of the 56 SNPs in SNP Set-2 had positive predictive performance. Utilizing the data from 61 cases within 69^*th*^-88^*th*^ range and 61 age-matching controls, AUC range was approximately the same (0.61-0.66). Similarly, only 19 SNPs in SNP Set-2 had positive predictive importance. These results further extended the robustness of our findings in both discovery and replication cohorts and demonstrated the specificity of the high predictive performance of SNP Set-2, which is derived from the cases in the ClinSeq® discovery cohort within 89^*th*^-99^*th*^ percentile CAC score range, to the advanced CAC phenotype.

### Identifying a subset of SNPs in SNP Set-2 leading to optimal predictive performance in both cohorts and function and disease enrichment analysis

Table 3 shows the list of 21 SNPs in SNP Set-2 generated optimal predictive performance in ClinSeq® and FHS cohorts. Using the genotypes of these 21 SNPs, we trained neural network models of 400 distinct topologies with ClinSeq® data and tested each topology with the FHS data. As shown in Figure 5A, we obtained 36 model topologies with AUC values ranging between 0.80-0.85 with empirically derived p-values of less than 0.05, thereby utilizing a different machine learning approach to replicate the collective predictive ability of these SNPs in the FHS replication cohort. This result demonstrates the stable and consistent features of these 21 SNPs in predicting advanced CAC independent of the classifier strategy employed. The optimal neural network topologies have 9-20 nodes in their first hidden layers and 6-20 nodes in their slightly less complex second hidden layers (Figure 5B).

**Table 3.**
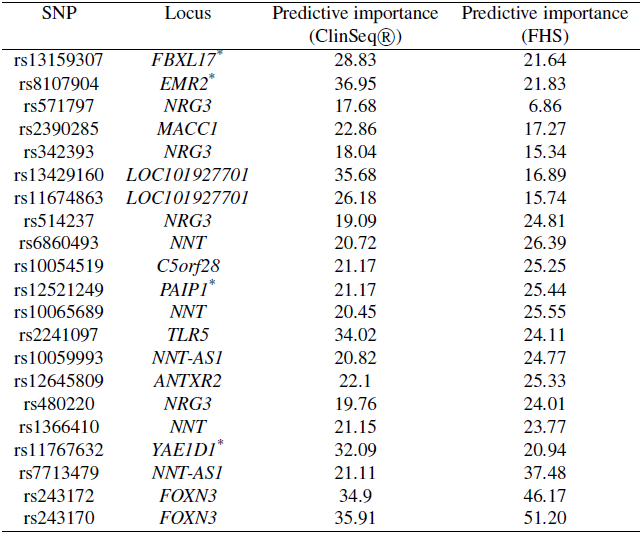
Predictive importance values of the set of SNPs that generate optimal predictive performance in both cohorts. Nearest genes are listed for intergenic SNPs (marked with asterisk). Predictive importance values of 12 of the 21 SNPs in the two cohorts are within 30% of each other (difference divided by the maximum value). In terms of predictive importance, five of the top 11 SNP predictors (with 65% of the cumulative predictive importance) are common, whereas nine of the top 14 SNP predictors (with 76% of the cumulative predictive importance) overlap between two cohorts.

**Figure 5.**
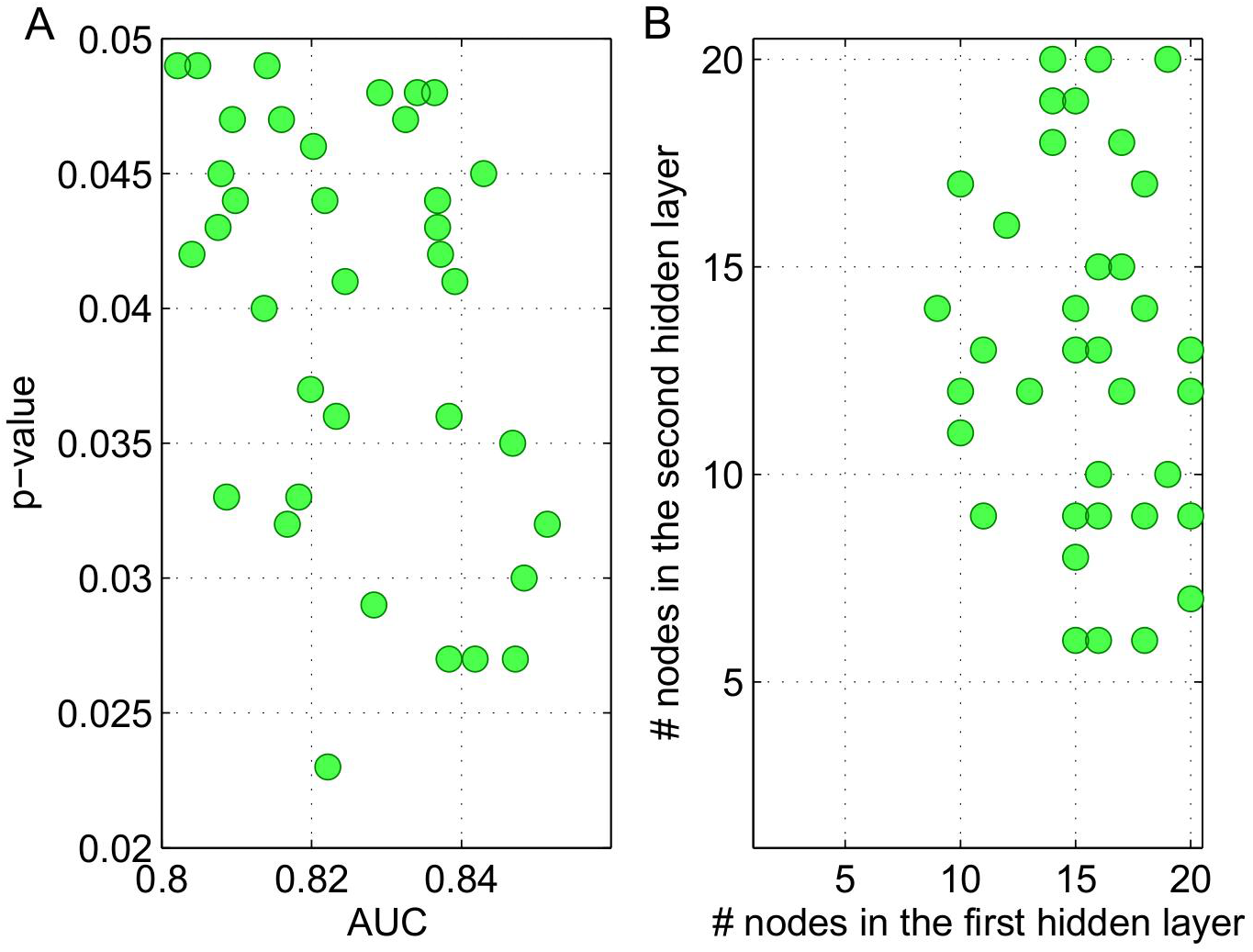
Properties of 36 optimal neural network models trained with data from the discovery cohort and tested with data from the replication cohort. A) Median AUC value for each network topology (ranging between 0.8021 and 0.8515) and the corresponding p-values. AUC distributions (one AUC distribution with 100 values per topology) were non-normal based on the Anderson-Darling tests Stephens (1974). Third quartile AUC values among the different network topologies ranged between 0.8503 and 0.9074. B) The number of nodes in the two hidden layers for each of the 36 optimal neural network topologies.

We identified a total of 13 genes that included the 21 SNPs leading to optimal predictive performance in both cohorts. Using GeneMANIA, we derived a network that included this group of 13 genes in addition to the 18 genes known to be linked to the first group based on coexpression and genetic interaction data from the literature (Warde-Farley et al., 2010). Figure 6 shows this network, whereas the abbreviated gene symbols and the corresponding gene names are listed in Table S10. The proteins coded by the genes in the network have a wide range of roles. 12 of them are either a transcription factor or an enzyme, one is a translational regulator, and two are transmembrane receptors.

**Figure 6.**
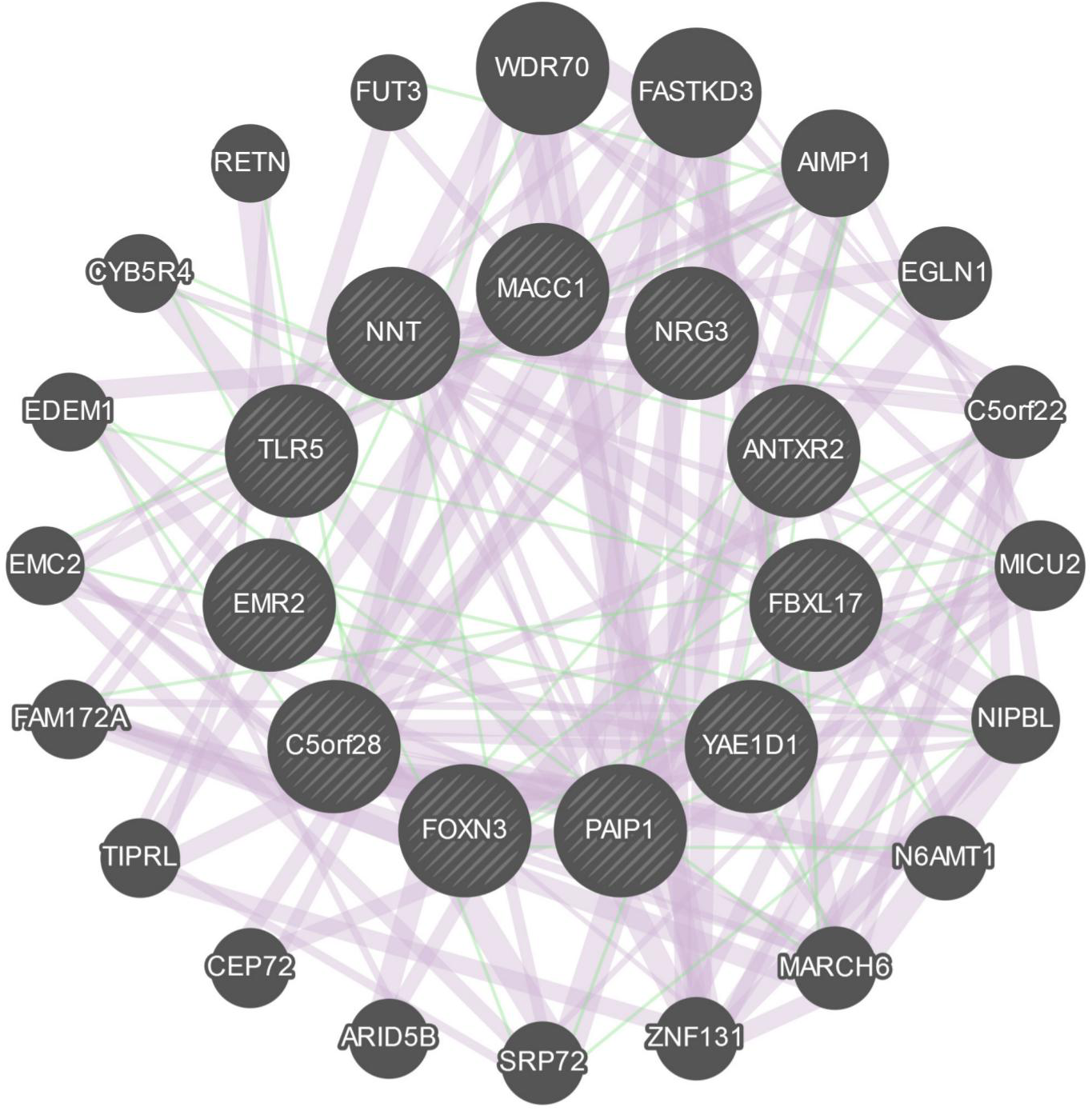
Network derived from GeneMANIA based on 244 studies in humans. The connections in pink are derived from gene coexpression data, whereas the connections in green are derived from genetic interaction data from the literature. The inner circle is composed of genes on which the subset of SNPs in SNP Set-2 leading to optimal performance in both cohorts are present, whereas the genes forming the outer circle are additional genes identified by GeneMANIA. The thicknesses of the links (or edges) between the genes are proportional to the interaction strengths, whereas the node size for each gene is proportional to the rank of the gene based on its importance (or gene score) within the network.

In order to identify whether our list of genes was enriched in any biological functions or processes associated with CAD, we used two bioinformatics resources, namely DAVID (Huang et al., 2009) and Ingenuity Pathway Analysis (IPA, Qiagen, Redwood City, CA, USA). Through their associations with blood magnesium levels (*NRG3*, *WDR70*, and *EMC2*), type-2 tumor necrosis factor receptors (*NRG3* and *ARID5B*), HDL cholesterol (*NRG3*, *MICU2*, *ARID5B*, and *FBXL17*), BMI (*AIMP1*, *MARCH6*, *FOXN3*, and *FAM172A*), CAD (*RETN*, *NNT*, *PAIP1*, and *MACC1*), respiratory function tests (*NRG3*, *EDEM1*, and *FAM172A*), and adiponectin (*NRG3* and *FBXL17*), 17 of the 31 genes in our network are associated with only one disease class, namely cardiovascular disease with a fold-enrichment of 1.9 and a p-value of 0.0025 (modified Fisher’s exact test) based on DAVID (Huang et al., 2009) and the Genetic Association Database.

Through mouse and rat models, six genes in our network have been previously associated with cardiovascular disease processes and risk factors. Several mouse models have linked *ARID5B* (a transcription factor involved in smooth muscle cell differentiation and proliferation) to obesity, differentiation of adipocytes, amount of white and adipose tissue, percentage body fat, and abnormal morphology of fat cells (Whitson et al., 2003; Rankinen et al., 2006; Yamakawa et al., 2008; Lahoud et al., 2001; Hata et al.,2013). Similarly, multiple mouse models (Rankinen et al., 2006; Xie et al., 2004; Xu et al., 2011; Zhang et al., 2010) showed that *CYB5R4* (involved in endoplasmic reticulum stress response pathway and glucose homeostasis) is associated with mass of adipose tissue, hypoinsulinemia, hyperglycemia, secretion of insulin, rate of oxidation of fatty acid, hyperlipidemia, timing of the onset of hyperglycemia, and diabetes. Similarly, using mouse model-based studies, *EGLN1* (involved in the regulation of angiogenesis, oxygen homeostasis, and response to nitric oxide) and its paralog EGLN3 have been linked to the necrosis of heart tissue, apoptosis of cardiomyocytes in infarcted mouse heart, stabilization of HIF1-alpha protein in left ventricle from mouse heart, functional recovery of heart, hepatic steatosis (fatty liver disease), angiectasis (abnormal dilation of blood vessels), and dilated cardiomyopathy (reduced ability of heart to pump blood due to enlarged and weakened left ventricle) (Hölscher¨ et al., 2011; Eckle et al., 2008; Takeda et al., 2006; Minamishima et al., 2009; Takeda et al., 2007). Through mouse and rat models, *RETN* (a biomarker for metabolic syndrome, atherosclerosis, and insulin-dependent diabetes, and a regulator of collagen metabolic process and smooth muscle cell migration) has been linked to insulin resistance, hyperinsulinemia, glucose intolerance, quantity of D-Glucose, quantity of circulating free fatty acid, LDLR, reactive oxygen species, and triglycerides (Satoh et al., 2004; Rajala et al., 2003; Steppan et al., 2001; Sato et al., 2005; Pravenec et al., 2003; Kim et al., 2004d). Several rat and mouse models showed that *TLR5* (a transmembrane receptor involved in inflammatory response, nitric oxide biosynthesis, and cellular response to lipopolysaccharide) is associated with obesity, hypertension, insulin resistance, autoimmune diabetes, cholesterol and triglyceride levels, systolic and diastolic blood pressure in systemic artery, and inflammation (Vijay-Kumar et al., 2010; Guo et al., 2006; Feuillet et al., 2006). Finally, *NRG3* serves as a ligand of the tyrosine kinase receptor ErbB4 that has been shown to affect the development of heart and the flow of blood in heart in multiple mouse models (Elenius and Paatero, 2008; Carpenter, 2003; Yarden and Sliwkowski, 2001; Tidcombe et al., 2003).

Table 4 shows the 22 cardiovascular disease related biological functions and phenotypes, which are identified by IPA based on Fisher’s exact test (p-value<0.01), enriched within our network of genes. Several of these functions and phenotypes are involved in biological processes associated with “vascular aging”, which is highly relevant to CAC, since aged vascular smooth muscle cells (VSMCs) are known to have less resistance against phenotypic modulations promoting vascular calcification (Shanahan, 2013). In fact, along with seven traditional risk factors (age, gender, total cholesterol, HDL cholesterol, systolic BP, smoking status, hypertension medication status), the Agatston CAC score is used as a parameter in quantifying “vascular age” in the MESA arterial age calculator (Dat, 2016b). Among our network genes previously linked to biological processes related to “accelerated” arterial aging, TLR5 is a member of the TLR (toll-like receptor) family as an established mediator of atherosclerosis due to its role in immune response through the induction of inflammatory cytokines (Kim et al., 2016) along with *RETN*, *ARID5B*, *NIPBL*, *EGLN1*, and *CYB5R4* affecting the adipose tissue quantity, an important driver of vascular pathology (Berg and Scherer, 2005; Demer and Tintut, 2011).

**Table 4.** Enriched diseases and biological functions (in the network of genes derived from GeneMANIA) with p-values ranging between 1.0E-4 and 1.0E-2 as identified by IPA based on Fisher’s exact test. 51 additional enriched diseases and biological functions (statistically less significant) with p-values ranging between 1.0E-2 and 5.0E-2 are listed in Table S11.

| Category | Disease or Function | Genes | p-value |
| --- | --- | --- | --- |
| Connective Tissue Development and Function | Quantity of adipose tissue | *ARID5B, CYB5R4, RETN, TLR5* | 3.58E-4 |
| Connective Tissue Development and Function | Differentiation of adipocytes | *ARID5B, EGLN1, NIPBL, RETN* | 8.82E-4 |
| Cardiovascular Disease | Angiectasis of blood vessel | *EGLN1* | 9.87E-4 |
| Cardiovascular System Development and Function | Area of capillary vessel | *EGLN1* | 9.87E-4 |
| Hematological System Development and Function | Cell division of peripheral blood lymphocytes | *AIMP1* | 9.87E-4 |
| Cardiovascular Disease, Endocrine System Disorders, Metabolic Disease | Susceptibility to insulin resistance-related hypertension | *RETN* | 9.87E-4 |
| Cardiac Necrosis, Cell Death and Survival | Cell death of heart tissue | *EGLN1* | 1.97E-3 |
| Cellular Movement | Migration of connective tissue cells | *AIMP1, ARID5B, RETN* | 2.14E-3 |
| Carbohydrate Metabolism, Cellular Function and Maintenance | Homeostasis of D-glucose | *CYB5R4, RETN, TLR5* | 2.46E-3 |
| Nucleic Acid Metabolism | Conversion of NAD+ | *NNT* | 2.96E-3 |
| Cardiovascular System Development and Function | Tethering of endothelial cell lines | *FUT3* | 2.96E-3 |
| Cellular Compromise, Inflammatory Response | Degranulation of beta islet cells | *CYB5R4* | 3.94E-3 |
| Cardiovascular System Development and Function | Density of blood vessel tissue | *AIMP1* | 3.94E-3 |
| Endocrine System Disorders, Hematological Disease Metabolic Disease | Onset of hyperglycemia | *CYB5R4* | 3.94E-3 |
| Carbohydrate Metabolism | Tolerance of D-glucose | *CYB5R4* | 4.93E-3 |
| Cardiovascular System Development and Function | Angiogenesis of heart | *EGLN1* | 5.91E-3 |
| Cardiovascular System Development and Function | Density of blood vessel | *AIMP1, EGLN1* | 5.96E-3 |
| Immune Cell Trafficking, Inflammatory Response Hematological System Development and Function | Adhesion of neutrophils | *ADGRE2 (EMR2), TLR5* | 7.52E-3 |
| Endocrine System Development and Function Hepatic System Development and Function | Insulin sensitivity of liver | *RETN* | 7.87E-3 |
| Nucleic Acid Metabolism | Metabolism of NADPH | *CYB5R4* | 7.87E-3 |
| Connective Tissue Development and Function | Quantity of visceral fat | *RETN* | 8.85E-3 |
| Carbohydrate Metabolism | Binding of chondroitin sulfate | *ADGRE2 (EMR2)* | 9.83E-3 |

*MICU2* plays a critical role in Ca^2+^ homeostasis as the gatekeeper of mitochondrial Ca^2+^ uniporter (MCU) (Patron et al., 2014) that is responsible for Ca^2+^ uptake into mitochondrial matrix, whereas blocking MCU leads to suppression of ROS production in the mitochondria (Pei et al., 2016). The disruption of Ca^2+^ homeostasis is an essential element of metabolic diseases and has been previously linked to endoplasmic reticulum (ER) stress (Arruda and Hotamisligil, 2015). Two genes in our network (*EDEM1* and *MARCH6*), are involved in the endoplasmic-reticulum-associated protein degradation (ERAD) pathway that targets and degrades misfolded proteins under stress conditions in order to prevent their accumulation. The importance of ERAD in the heart has previously been established (Razeghi and Taegtmeyer, 2005; Wang and Robbins, 2006) especially for proper functioning of cardiomyocytes. In more recent studies, improving ERAD has been shown to preserve heart function and reduce cardiomyocyte death with mouse models of cardiac hypertrophy (Doroudgar et al., 2015) and MI (Belmont et al., 2010), respectively.

Through ROS activity, macrophages that contain lipid molecules (or foam cells) accumulate in the artery walls and promote atherosclerosis (Stocker and Keaney, 2004). *EMR2,* which is one of our network genes that promotes the release of inflammatory cytokines from macrophages, has been reported to be highly expressed in foamy macrophages handling lipid overload in atherosclerotic vessels (van Eijk et al., 2010). Excessive ROS generation (previously linked to vascular calcification (Johnson et al., 2006)) also leads to reduced levels of nitric oxide (NO) (Muzaffar et al., 2005), molecule with cardioprotective features. The reduced form of NADP (NADPH) is required for the synthesis of cholesterol (Lieberman et al., 2013) as a cofactor in all reduction reactions and also for the regeneration of reduced glutathione (GSH) (Gorrini et al., 2013) that providing protection against ROS activity (Murphy, 2012). Two of our network genes, NNT and *CYB5R4*, are involved in NADPH metabolism. Taken together, these findings show that several biological processes and risk factors previously linked to cardiovascular disease, and particularly to vascular aging, are enriched within the network we derived from the loci of SNPs that are highly predictive of advanced CAC.

## DISCUSSION

Understanding the drivers of accelerated CAD pathogenesis hold great potential for providing novel pathobiological insights into biological events, including inflammatory and immune responses (Björkegren et al., 2015; Libby et al., 2009; Hansson, 2005), beyond conventional mediators, such as the dysregulation of lipid metabolism or blood pressure (Björkegren et al., 2015; Roberts, 2014). A major goal in the cardiovascular disease field is identifying individuals who are at greatest risk of accelerated CAD pathogenesis. Recognizing that the utility of traditional risk factors (particularly those driven by age) is not sufficiently robust to identify all patient groups with accelerated CAD (Thanassoulis and Vasan, 2010), turning to genomic data and utilizing non-traditional statistical tools for building predictive models of CAD is a fairly recent avenue in biomedical research (Völzke et al., 2013). To this end, our study is an example of a machine learning-based predictive modeling approach that utilizes clinical and genotype data to identify a panel of SNPs providing improved predictive performance over traditional risk factors and a past GWAS-based panel in a replicable manner in two independent cohorts.

Recent literature suggests that the implementation of regression models using a log additive (or multiplicative) approach when integrating multiple SNPs together for making predictions (Yoo et al., 2015) is a potential pitfall in previous attempts to improve the risk prediction accuracy for complex diseases. Alternative modeling approaches that utilize SNPs while taking into account gene-gene and gene-environment effects are some of the promising potentials of “recursive partitioning methods” (Breiman, 2001; Ruczinski et al., 2003) including random forest models (Yoo et al., 2015). In our study, using random forests, we observed significantly improved predictive performance upon combining traditional risk factors with a past GWAS based SNP panel (SNP Set-1) in the discovery cohort as opposed to only using clinical data or SNP Set-1. On the other hand, in the replication cohort, combining clinical data with SNP Set-1 led to a slight decline in predictive performance compared to using only SNP Set-1, but resulted in a significant improvement as opposed to using only clinical data. Furthermore, we observed no predictive improvement in either cohort as we combined these clinical variables with the alternative set of SNPs (SNP Set-2) derived from the discovery cohort based on genotype-phenotype correlation. Taken together, our results are in accord with majority of the previous results in the literature since combining both layers of data have not generated a consistent improvement in our discovery and replication cohorts.

We note that in a previous predictive modeling study on CAC (McGeachie et al., 2009), authors have significantly improved the ability for predicting the presence of coronary calcium by combining clinical variables with 13 predictive SNPs from 13 different genes identified among 2882 candidate SNPs from 231 genes that were proposed by a group of MESA investigators. However, the data used in (McGeachie et al., 2009) came from a patient group with significantly different characteristics. Half of the patients in (McGeachie et al., 2009) were females, whereas our patients in both ClinSeq® and FHS cohorts were all males with much higher levels of coronary calcium. In fact, the CAC scores of our male case subjects were within 89^*th*^ to 99^*th*^ percentile range based on the Multi-Ethnic Study of Atherosclerosis (MESA) cohort (McClelland et al., 2006; Dat, 2015), whereas majority of the male data in (McGeachie et al., 2009) came from subjects with CAC scores within 60^*th*^-70^*th*^ percentile range based on the reported average age and the CAC score range. Hence, our case definition that is based on the presence of advanced coronary calcium, rather than its mere presence, in addition to the differences in the gender composition between cohorts, are plausible explanations for the discord between our study and (McGeachie et al., 2009) in terms of the changes in predictive performance upon combining clinical and genotype data.

In a recent review by Björkegren et al. (Björkegren et al., 2015), authors discuss the importance of nominally significant (p-value<0.05) SNPs that fail to reach genome-wide significance (p-value< 10^−8^) in terms of collectively explaining the genetic variability in CAD. The effectiveness of this approach has previously been shown in the context of the heritability of human height in (Gibson et al., 2010; Yang et al., 2010a). Nominally significant (also called “context-dependent”) SNPs show their impact on disease phenotypes only under certain conditions (Schadt and Björkegren, 2012), such as above a certain BMI threshold (LyssenkoMaas et al., 2008) or below some physical activity level (Rankinen et al., 2007). In (Björkegren et al., 2015), such variants are described as potential key drivers of CAD in later stages as opposed to GWAS significant loci that promote early development of CAD. Based on this argument (also supported by a recent study (Roberts, 2014)) early CAD development is driven mainly by genetics rather than environmental factors, as opposed to the context-dependent variants that drive later stages of CAD and are typically unable to reach genome-wide significance. However, as demonstrated in past studies (Björkegren et al., 2015; Schadt and Björkegren, 2012), it’s possible to utilize context-dependent variants for building predictive disease models by integrating multiple layers of omics data with clinical variables. Our study is an example of such an integrative approach and the results in Tables S6 and S8 demonstrate the emergence of several SNPs previously associated with several CAD risk factors (mostly at nominal significance) in driving advanced CAC levels as demonstrated by the cumulative predictive scores attached to each risk factor. In addition, the associations between the predictive SNPs in SNP Set-2 with CAD-related phenotypes (Table S9) identified in previous GWAS were all nominally significant contrary to the predictive SNPs from SNP Set-1 derived from past GWAS on CAC, many of which reached genome-wide significance previously in GWAS on CAD-related phenotypes (Table S7). In (Björkegren et al., 2015), the recommended approach for predicting CAD and related phenotypes (especially beyond early disease stages) is dividing case subjects into subcategories based on the level of disease measured by imaging or histological measures (measured CAC scores in our study) to identify subphenotype-specific integrative models. We implement a similar approach in our predictive modeling study by just focusing on case subjects within the 89^*th*^-99^*th*^ percentile CAC score range and age-matching controls. The replication of the highly predictive loci identified from the ClinSeq® discovery cohort in the FHS cohort and the fact that we observe enrichment of several biological processes previously linked to cardiovascular disease at the network level demonstrates the effectiveness of our machine learning based approach.

## CONCLUSIONS

In this study, we used a combination of clinical and genotype data for predictive modeling of advanced coronary calcium. Our models demonstrated the limited predictive capabilities of traditional risk factors and a past GWAS-based SNP panel, whereas an alternative SNP set, with approximately the same size as the GWAS-based panel, produced higher predictive performance in a discovery cohort from ClinSeq® study and in a replication cohort from FHS. 75% of the SNPs in this alternative set have previously been associated with a total of 18 risk factors (a total of 88 associations), whereas 18% of them have reached nominal significance levels in a previous GWAS on mitral annular and aortic valve calcium that suggested potentially strong susceptibility to CAD as well as coronary calcium among our subjects with advanced CAC. Upon identifying a subset of 21 SNPs from this alternative set that led to optimal predictive performance in both cohorts, we developed neural network models trained with the ClinSeq® genotype data and tested with the FHS genotype data and obtained high predictive accuracy values (AUC>0.8) under a wide range of network topologies, thereby replicating the collective predictive ability of these SNPs in FHS and identifying several potential susceptibility loci for advanced CAD pathogenesis. At the gene network level, several biological processes previously linked to cardiovascular disease, including differentiation of adipocytes, were found to be enriched among these loci.

A potential extension of our modeling study is the expansion of the panel of SNPs that are highly predictive of advanced coronary calcium levels around their loci for building more comprehensive models. Subsequently, we can assess these loci as predictors of rapid CAC progression and early onset of MI with longitudinal data in independent cohorts, especially for cases poorly predicted by traditional risk factors. To conclude, our study on CAC, a cardiovascular disease phenotype and a predictive marker of future cardiac events, demonstrates the limited capability of the GWAS-based set of markers in predicting advanced CAC, while illustrating the potential of combining multiple machine learning methods as informative and accurate diagnostic tools. Our results also suggest that utilizing markers specific to a particular range of coronary calcium, rather than its complete spectrum previously studied in past GWAS, can be an effective approach for building accurate predictive models for personalized medicine efforts that require disease-level specific risk prediction and prevention.

## AUTHOR’S CONTRIBUTIONS

Conceived the study: CO, SKS, ARD, YPF, CJO, GHG. Developed the methodology, performed the statistical modeling and analysis: CO. Wrote the paper: CO, SKS, ARD, YPF, CJO, GHG. All authors read and approved the final manuscript.

## ACKNOWLEDGMENTS

The authors gratefully acknowledge the Intramural Program of the National Human Genome Research Institute of the National Institutes of Health for funding this research. We also gratefully acknowledge Leslie Biesecker for contribution of ClinSeq® data, which was funded by NIH grants HG200359 08 and HG200387 03. The views expressed in this manuscript are those of the authors and do not necessarily represent the views of the National Heart, Lung, and Blood Institute; National Human Genome Research Institute; the National Institutes of Health; or the U. S. Department of Health and Human Services.

